# The SARS-CoV-2 infection in Thailand: analysis of spike variants complemented by protein structure insights

**DOI:** 10.1101/2022.01.01.474713

**Authors:** Sirawit Ittisoponpisan, Shalip Yahangkiakan, Michael J E Sternberg, Alessia David

## Abstract

Thailand was the first country outside China to officially report COVID-19 cases. Despite the strict regulations for international arrivals, up until February 2021, Thailand had been hit by two major outbreaks. With a large number of SARS-CoV-2 sequences collected from patients, the effects of many genetic variations, especially those unique to Thai strains, are yet to be elucidated. In this study, we analysed 439,197 sequences of the SARS-CoV-2 spike protein collected from NCBI and GISAID databases. 595 sequences were from Thailand and contained 52 variants, of which 6 had not been observed outside Thailand (p.T51N, p.P57T, p.I68R, p.S205T, p.K278T, p.G832C). These variants were not predicted to be of concern. We demonstrate that the p.D614G, although already present during the first Thai outbreak, became the prevalent strain during the second outbreak, similarly to what was described in other countries. Moreover, we show that the most common variants detected in Thailand (p.A829T, p.S459F and p.S939F) do not appear to cause any major structural change to the spike trimer or the spike-ACE2 interaction. Among the variants identified in Thailand was p.N501T. This variant, which involves an asparagine critical for spike-ACE2 binding, was not predicted to increase SARS-CoV-2 binding, thus in contrast to the variant of global concern p.N501Y. In conclusion, novel variants identified in Thailand are unlikely to increase the fitness of SARS-CoV-2. The insights obtained from this study could aid SARS-CoV-2 variants prioritisations and help molecular biologists and virologists working on strain surveillance.

## Introduction

In January 2020, Thailand became the first country outside China to officially report a COVID-19 case [1]. The local transmissions later developed into the first wave in March 2020, resulting in a declaration of a state of emergency, followed by a lockdown nationwide [2]. As part of the new regulations aiming to contain the outbreak, many international flights were banned, and individuals arriving in Thailand from overseas were required to stay in a 14-day state quarantine for monitoring before being released [3], rendering Thailand into virtual isolation. The lockdown was extensively lifted in July 2020 as no local transmission had been reported for over a month, flattening the infection tally to about 3,000 cases [4, 5]. However, citizens were still strongly advised to follow the “new normal” lifestyle of wearing face masks and maintaining social distancing in public areas.

Despite the government’s strict travel regulations in an effort to control cross-border transmission, there had been reports of local transmission cases due to illegal border crossing and not complying with the 14 day-quarantine measure, importing new strains from abroad [6–8]. In December 2020, the second wave of infection occurred among Burmese workers in a seafood market in Samut Sakhon province located to the southwest of Bangkok. It later quickly spread to the surrounding provinces [9]. As of February 2021, there had been reports of over 20,000 COVID-19 cases in Thailand and over a hundred million cases worldwide [10].

The rapid growth in COVID-19 cases worldwide and the global sequencing efforts resulted in a large number of viral sequences being collected from patients. Many of these have been made freely available through two major databases: NCBI (https://www.ncbi.nlm.nih.gov/sars-cov-2/) and GISAID (https://www.gisaid.org/) [11]. At the time of this study, over 600,000 SARS-CoV-2 sequences were available for analysis.

The spike protein is one of the most widely studied proteins of SARS-CoV-2 as it interacts with the human ACE2 (hACE2), thus facilitating viral cell entry [12, 13]. Analysis of the 3D structure of the spike protein, alone and bound to hACE2, can help researchers understand SARS-CoV-2 evolution and transmission and could guide therapeutic research [14]. To date, most of the analyses have been performed on variants of global concern identified in Europe, America and Africa, where the infections were severe and the fatality rates high [15–17]. However, studies on the effect of variants identified in Thailand are limited [18–21].

In this study, we aim to provide structural insights into SARS-CoV-2 spike variants identified in Thailand to obtain insights into SARS-CoV-2 Thai outbreaks and identify variants of interest which may require additional investigations.

## Methods

### SARS-CoV2 spike protein sequences

The spike protein sequences were retrieved on 7 February 2021 from two databases: NCBI (https://www.ncbi.nlm.nih.gov/sars-cov-2/) and GISAID (https://www.gisaid.org/) [11]. All sequences that were 1,273 amino acids long, which is the same length as the original and canonical Wuhan reference sequence (YP_009724390.1), were selected from these databases. As a result, 53,056 spike protein sequences from NCBI (including the reference YP_009724390.1) and 386,176 sequences from GISAID were retrieved. The sequences were then further screened by comparing them with the Wuhan reference sequence (YP_009724390.1). Any sequence that resulted in less than 98% identity or more than three consecutive amino acid variants were removed from the dataset. This is to eliminate the possibility of the sequence containing insertion/deletion mutations. From these criteria, six sequences from NCBI and 29 sequences from GISAID were removed. The final dataset consisted of 53,050 sequences from NCBI and 386,147 from GISAID (439,197 total).

### Phylogenetic analysis

The phylogenetic tree was generated using the Nextstrain web server [22]. The tree was created based on the whole genome alignment of SARS-CoV-2 strains available on the GISAID database and was customised to cover only strains collected in Thailand from 20 December 2019 to 7 February 2021 [23]. As Nextrain only displays complete strains on a fixed timeline, some variants could not be mapped directly onto this tree due to incomplete strain information (i.e., the strain might be partially sequenced, or the date of collection was unclear). Therefore, the locations of some of these variants were estimated based on their clade information and co-existing variants on the strain.

### Structural analysis

The following 3D coordinate files were extracted from the ProteinDataBank and used for variant analysis: 6XR8 (resolution: 2.90 Å, released: 2020-07-22) for the complete structure of the SARS-CoV-2 spike protein complex (at inactive conformation) and 6M0J (resolution: 2.45 Å, released: 2020-03-18) for the spike-hACE2 complex. The accessible surface area (ASA) calculation was performed using DSSP [24]. For each amino acid, the RSA was calculated using the formula RSA = ASA/maxASA. The maxASA for each amino acid is defined according to Rost and Sander [25]. A residue was regarded as “surface residue” when its relative solvent accessibility (RSA) was ≥ 9%, otherwise “buried”. A residue was defined as an “interface” if within 4 Angstrom distance from any other residues of a different chain [26, 27]. In this study, interface residue calculation was performed on the trimer spike protein complex (PDB: 6XR8) and the spike-hACE2 complex (PDB: 6M0J). Full details of surface and interface residues are provided in Supplementary File 1.

Variants occurring in the SARS-CoV-2 spike were identified by comparing 439,196 sequences in our dataset against the Wuhan SARS-CoV-2 spike reference sequence (YP_009724390). The in silico mutagenesis was performed using a modified version of the Missense 3D algorithm [28] to account for amino acid substitutions at interface residues.

## Results

### Phylogenetic analysis of variants found in Thailand

A total of 439,197 SARS-CoV-2 spike sequences were collected and analysed. 595 sequences were from Thailand and contained 52 spike variants. Forty-one of these variants could be mapped onto a phylogenetic tree (Figure 1). Two major clusters of strains from the first and second SARS-CoV-2 Thai outbreaks were identified. The first outbreak contained variants from seven clades of SARS-CoV-2 strains. Although the variant of global concern p.D614G was already detected in many strains since March 2020, it became the prevalent strain in the second outbreak (clade GH, Figure 1).

**Figure 1:**
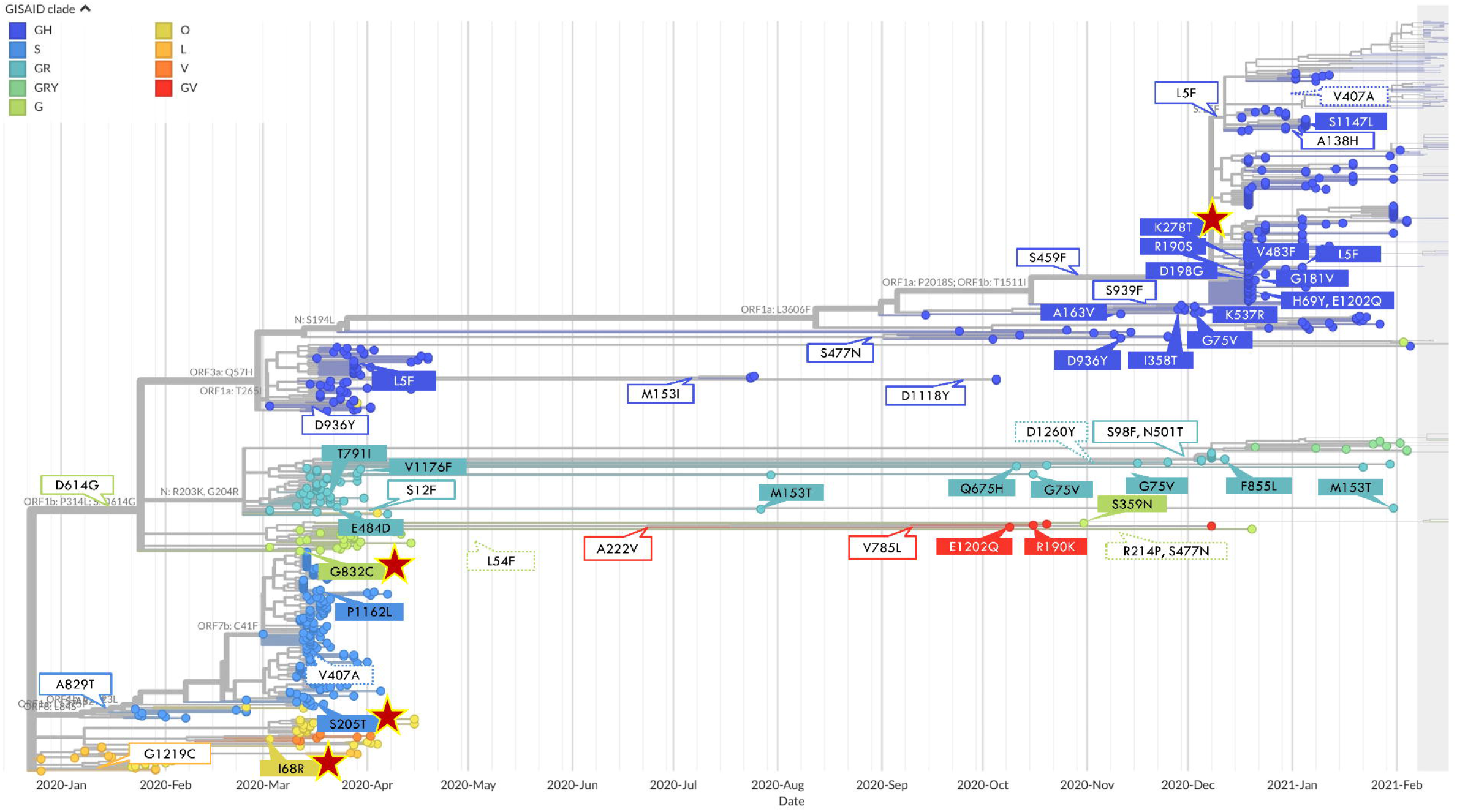
Phylogenetic analysis of SARS-CoV-2 strains collected in Thailand from December 2019 to February 2021. In the phylogenetic tree, each node represents a strain. White boxes with colour borders representbranches where variants evolved and are shared among descendants. Colour boxes indicate newly evolved variants found on each particular leaf node. Variants that cannot be mapped directly onto this tree due to incomplete strain information are estimated and represented as white boxes with dotted borders. Variants that were unique to Thai strains are indicated by the red asterisk (⋆).

Six SARS-CoV-2 spike variants, p.T51N, p.P57T, p.I68R, p.S205T, p.K278T, p.G832C, were unique to Thai strains and not identified in genetic sequences from other countries. Notably, three variants (p.I68R, p.S205T, and p.G832C) were found during the first outbreak, and one (p.K278T) was found during the second outbreak. The other two Thailand-specific variants (p.T51N and p.P57T), which co-localized in one genomic sequence, could not be mapped onto the phylogenetic tree (ID: 708814, clade: GH, collected on 9 October 2020). This strain had a total of 11 variants: eight of them – p.V47A, p.T51N, p.Q52K, p.F55L, p.P57T, p.V70L, p.S71P, and p.A688P - were not shared by any other Thai strains. When the whole genome of this record was used in a BLAST search against SARS-CoV-2 records on the GISAID web server, similar strains were found mainly in samples from Saudi Arabia (data not shown). This suggests that this strain was not a result of local transmission but may have been introduced into Thailand from overseas and later mutated to contain the two unique variants p.T51N and p.P57T.

### Analysis of variants found in Thailand

We first analysed the distribution of all variants identified in the spike protein by using the entire dataset of 439,197 SARS-CoV-2 spike sequences. On average, each spike sequence had 2.7 variants (min: 0, max: 23, median: 3, STD: 1.17) (see Supplementary File 2). Variants were distributed ubiquitously on the spike amino acid sequence, with the exception of 51 residues which did not contain any variants. These residues are part of the receptor-binding (RBD) and Corona S2 superfamily (S2) domains (Supplementary Figure 1 and Supplementary File 3-4).

We next focused on the six variants uniquely identified in Thailand (see Table 1 and Figure 2). No structural changes to the spike trimer and the spike-hACE2 interaction were identified with the exception of p.S205T, which was predicted to reduce the structural stability of the spike protein due to drastic reduction of a surface cavity. Cavity alteration has been shown to affect protein stability [14–16]. Hence, it is likely that this variant is deleterious to the virus. Recently, p.I68R was discovered in a mutagenesis study of SARS-CoV-2 strains in mice and reported to be associated with the virus’s ability to escape neutralising antibodies [29]. However, whether this variant enhances the viral escape mechanism in humans is yet to be elucidated. Despite no structural damage found from p.K278T, the extra h-bond formed between the main chain of Thr278 and the side chain of Thr286 was due to the side chain repacking of Thr286 rather than the mutation from Lys278 to Thr278. Nevertheless, the fact that all of the six Thailand-specific variants were rare (found in < 1% of the samples collected) could imply that they are unlikely to enhance the virus’s transmissibility. We next examined additional variants identified in Thailand (the full structural analysis of the 52 spike variants found in Thailand is provided in Supplementary Table 1). p.D614G was identified in 48.7% of all sequences analysed. This variant was predicted to cause the loss of inter-chains H-bonds between aspartic acid 614 and lysines 835 and 854 in the spike trimer, which could affect the spike trimer packing. A study by Yurkovetskiy et al. using cryo-electron microscopy, indeed, confirms that p.D614G disrupts an interprotomer contact, thus facilitating the spike “open conformation” [16, 30]. This variant is estimated to increase the transmissibility of SARS-CoV-2 by 20% [31], thus explaining why, although identified in few samples in March 2020 (first Thai outbreak) before a strict lockdown measure was enforced, it had become the dominant strain by February 2021 (second Thai outbreak), similarly to what was observed in other South East Asian countries [32, 33]. The higher transmissibility of p.D614G can also explain the higher number of SARS-CoV-2 positive cases in the second compared to the first Thai outbreak.

**Table 1:**
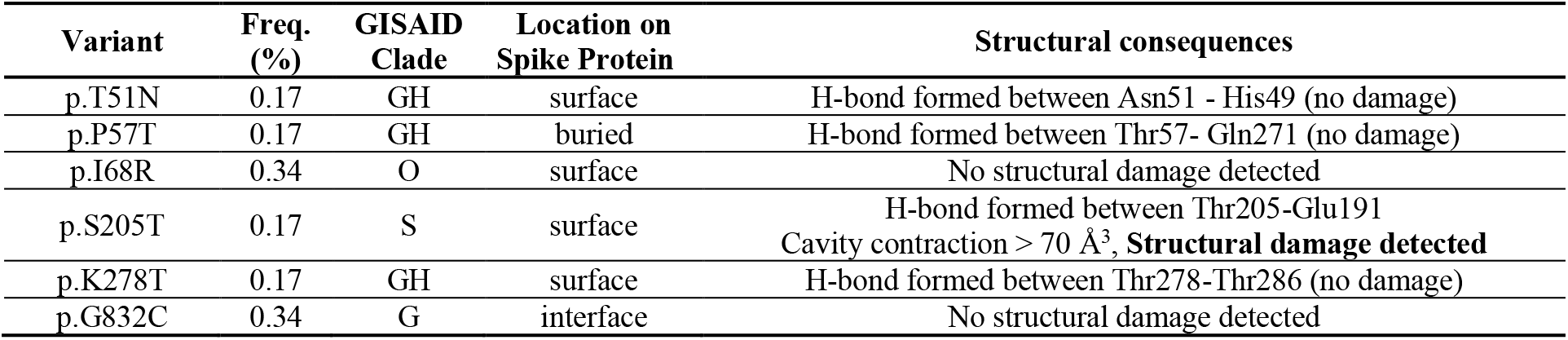
Analysis of six variants found specifically in Thailand.

**Figure 2:**
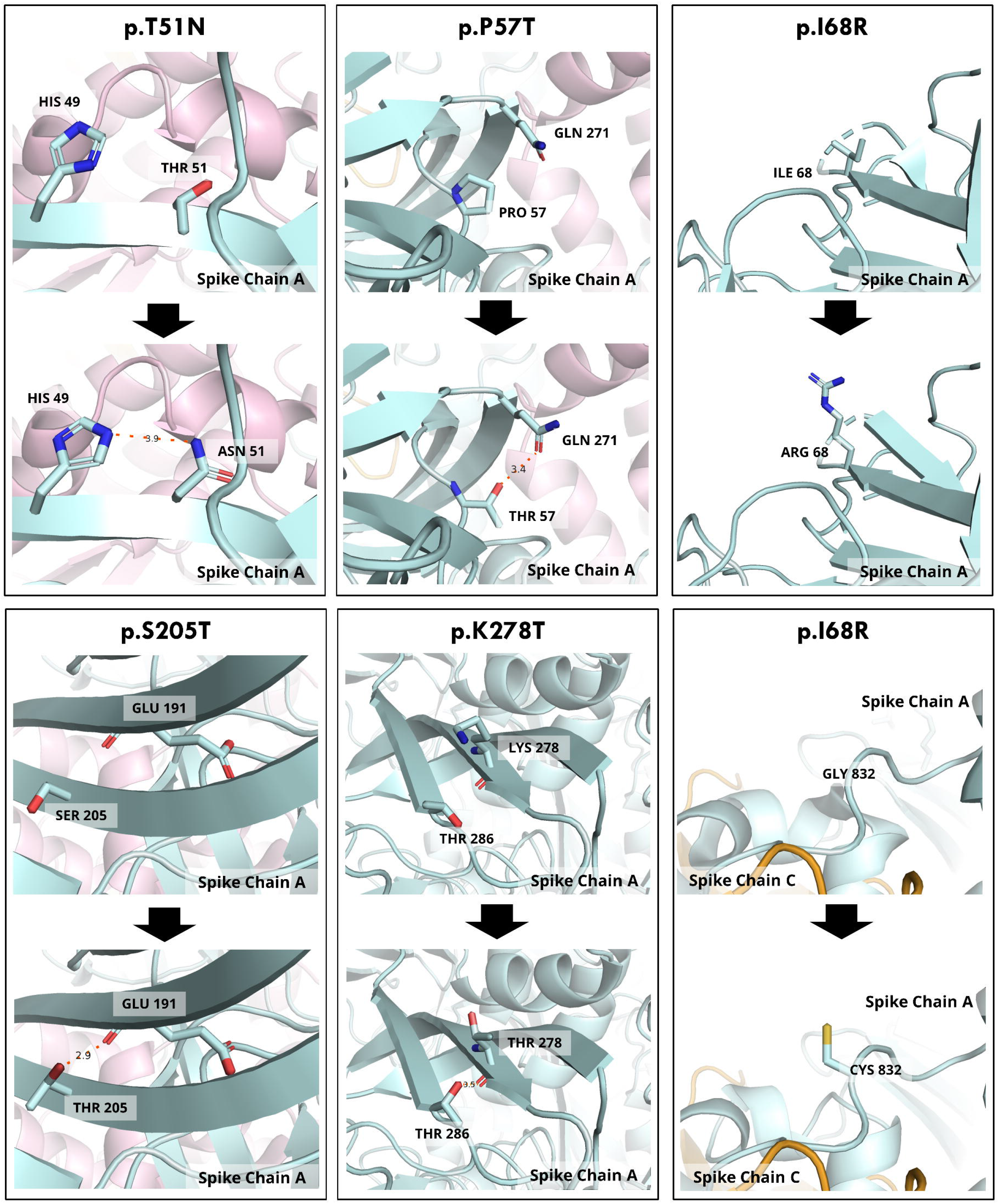
Structural analysis of Thai unique variants. In each panel, the wild type structure is presented on top and the predicted mutant on the bottom. The PDB structure used in this analysis was 6XR8. Chain A, B, and C are represented in pale blue, light pink, and yellow, respectively. All variants were simulated on Chain A. H-bonds are shown as orange dashed lines.

The other most common variants identified in Thailand were p.A829T (39.2% of all sequences), p.S459F (20.2%), p.L5F (4.7%), p.S939F (2.0%) and p. N501T (1.5%). p.A829T was the prevalent SARS-CoV-2 variant during the first Thai outbreak, but it was no longer detected in the second outbreak. No clear structural damage was identified by our analysis. Variant p.S459F was predicted to disrupt an H-bond formed within surface residues (an H-bond linking Arg457 and Ser459 was disrupted when the variant was simulated on PDB 6XR8, while a disruption of an H-bond formed between two consecutive residues Ser459 and Asn460 was detected when tested on PDB 4M0J), which is likely to be compensated by the interaction with nearby water molecules and therefore not structurally destabilising. Variant p.L5F was a newly emerging variant in the second Thai outbreak. It is located near the N-terminal of the spike protein. Unfortunately, this residue was not covered by the available 3D coordinates and no structural analysis could be performed. Variant p.S939F was not predicted to affect the structure of the spike trimer or its interaction with ACE2. Accordingly, a previous study suggested that this variant is not likely to enhance SARS-CoV-2 infectivity [34].

Variant p. N501T was the only amino acid substitution occurring at the interface between spike and the hACE2. Asparagine 501 is a critical residue involved in the stabilization of lysine 353, one of the two critical lysines involved in spike-hACE2 interaction [35]. Indeed, asparagine 501 harbors the variant of concern p.N501Y, which has been proposed to enhance spike-hACE2 affinity through the introduction of new hydrogen bonds between tyrosine 501 on the spike protein and residues on the hACE2 [17, 36, 37]. We analysed the structural consequences of p.N501T and did not observe major structural changes. Indeed, an in vitro study showed this variant may cause a reduction in spike-hACE2 binding affinity [35].

## Discussion

At the time of this study, limited information was available on variants identified in Thailand during the two major SARS-CoV-2 outbreaks. Our phylogenetic analysis shows how the variant of global concern p.D614G already present in March 2020 became the dominant strain during the second Thai outbreak in late December 2020 - February 2021. This variant was detected in the majority of the strains collected from the seafood market [38, 39]. Found predominantly in Europe, this spike protein variant has been shown to have a higher transmissibility rate compared to the original Wuhan strain [31, 40]. The screening of variants present in Thailand also identified variant p.N501T. Aspargine 501 is a critical residue for spike-hACE2, and its substitution to tyrosine is a variant of concern. However, our analysis did not suggest that the asparagine to threonine substitution may cause any major structural change. Our results are supported by the in vitro analysis performed by Shang et al. ^(2020b)^, which did not show an increased spike-hACE2 binding affinity.

Interestingly, despite analysing nearly 500,000 sequences, some residues in the SARS-CoV-2 spike protein did not contain any variant. Many of these highly conserved residues are clustered in the receptor-binding domain, which binds to the human ACE2 and triggers host cell invasion [12, 13, 35], and the Coronavirus S2 domain, which mediates viral cell membrane fusion [41]. The lack of variants in these regions may suggest biological importance of those particular residues on protein folding and/or interaction. As genetic variations are usually maintained by natural selection, any alteration on these highly conserved residues could be deleterious to the survival of the virus. Conserved domains and residues could therefore serve as biomarkers [42] or promising drug targets against SARS-CoV-2 [41, 43].

One limitation in this study is that the Thai sequences available on NCBI and GISAID were from two sources: the state quarantines and domestic hospitals/research institutions. The strains collected in the state quarantine zones are likely imported in Thailand and less likely to cause local transmission. Unfortunately, the available data did not allow to distinguish the variant source. Hence, the number of variants found in Thailand in this study could be an overestimation of the variants that actually caused local transmissions. Nevertheless, it is always important to keep tracking of possible new strains as the mutation rate is very high in single-stranded RNA viruses compared to DNA viruses [44]. The high mutation rate poses a great challenge for developing vaccines [45], highlighting why incorporating conserved residue information into structural analyses could be essential for discovering other alternative measures for COVID-19 diagnosis, treatment, and prevention.

Another limitation is that at the time of this study only the 3D coordinates of the spike trimer and the spike-hACE2 interaction were available. Recently, the spike protein was reported to bind to other human proteins, such as CD209 [46], HAVCR1 [47], and NRP1 [48, 49], and other mammal proteins [50]. Therefore, in this study, the number of interface residues could have been underestimated, and it is possible that in the future, additional variants will be classified according to their effect on additional virus-host interaction. It is well established that interface residues are hot spots for disease variants [51–53]. The p.N501Y is an example that shows how viral variants occurring at protein interface could increase viral transmissibility and highlights that any SARS-CoV-2 variant occurring at the spike trimer interface or spike-hACE2 interface should be carefully analysed.

## Conclusions

In this study, we explored the structural effect of SARS-CoV-2 spike variants identified in Thailand, highlighting how the second Thai outbreak was likely caused by the variant of global concern p.D614G. Additionally, we highlighted highly conserved residues on the spike protein, which could have implications for the development of biomarkers or drug targets. The insights obtained from this study could aid SARS-CoV-2 variants prioritisations and help molecular biologists and virologists working on strain surveillance

## Declarations

### Availability of data and materials

The spike protein sequences from NCBI are available at (https://www.ncbi.nlm.nih.gov/sars-cov-2/). The GISAID spike protein sequences are available to registered users at (https://www.gisaid.org/). The three-dimensional structures for the Sars-Co-V2 are available from the ProteinDataBank at https://www.rcsb.org. All data generated or analysed during this study are included in this published article and its supplementary information files.

### Authors’ Contribution

SI conceptualised and designed the main methodology. SY performed data curation and investigation. SI conducted the analysis and wrote the manuscript. AD reviewed the original draft. All authors have read and agreed to the published version of the manuscript.

## References

[1] WHO. WHO Timeline COVID-19 https://www.who.int/news/item/27-04-2020-who-timeline---covid-19 (accessed Apr 23, 2021).

[2] WHO Thailand. WHO Thailand situation report - 41 https://www.who.int/docs/default-source/searo/thailand/2020-04-3-tha-sitrep-41-covid19-final.pdf?sfvrsn=9e14aebc_0.pdf (accessed Apr 23, 2021).

[3] Department of Disease Control. Measures to control COVID-19 Version 5 updated on 2 April 2020 https://ddc.moph.go.th/viralpneumonia/eng/file/main/Measures_for_traverlers_from_aboard_v5.pdf (accessed Apr 23, 2021).

[4] Department of Disease Control. Special Announcement of COVID-19 On 1 July 2020 https://ddc.moph.go.th/viralpneumonia/eng/file/news/news_no157_010763.pdf (accessed Apr 23, 2021).

[5] WHO Thailand. WHO Thailand situation report - 95 https://www.who.int/docs/default-source/searo/thailand/2020-07-06-tha-sitrep-95-covid19.pdf?sfvrsn=272b28fd_2 (accessed Apr 23, 2021).

[6] Department of Disease Control. Thailand situation update on 22 November 2020 https://ddc.moph.go.th/viralpneumonia/eng/file/situation/situation-no322-221163.pdf (accessed Apr 24, 2021).

[7] Department of Disease Control. Thailand situation update on 28 November 2020 https://ddc.moph.go.th/viralpneumonia/eng/file/situation/situation-no328-281163.pdf (accessed Apr 24, 2021).

[8] Department of Disease Control. Thailand situation update on 3 December 2020 https://ddc.moph.go.th/viralpneumonia/eng/file/situation/situation-no333-031263.pdf (accessed Apr 24, 2021).

[9] WHO Thailand. WHO Thailand situation report - 102 https://cdn.who.int/media/docs/default-source/searo/thailand/2020_12_21_tha-sitrep-102-covid19_r02.pdf?sfvrsn=7e62038c_7&Status=Master (accessed Apr 24, 2021).

[10] Worldometers.info. COVID-19 Coronavirus Pandemix https://www.worldometers.info/coronavirus/ (accessed Apr 2, 2021).

[11] Bogner, P.; Capua, I.; Cox, N. J.; Lipman, D. J. A Global Initiative on Sharing Avian Flu Data [1]. Nature. Nature Publishing Group August 31, 2006, p 981. https://doi.org/10.1038/442981a.

[12] Lan, J.; Ge, J.; Yu, J.; Shan, S.; Zhou, H.; Fan, S.; Zhang, Q.; Shi, X.; Wang, Q.; Zhang, L.; et al. Structure of the SARS-CoV-2 Spike Receptor-Binding Domain Bound to the ACE2 Receptor. Nature, 2020, 581 (7807), 215–220. https://doi.org/10.1038/s41586-020-2180-5.

[13] Shang, J.; Wan, Y.; Luo, C.; Ye, G.; Geng, Q.; Auerbach, A.; Li, F. Cell Entry Mechanisms of SARS-CoV-2. Proc. Natl. Acad. Sci. U. S. A., 2020, 117 (21), 11727–11734. https://doi.org/10.1073/pnas.2003138117.

[14] Waman, V. P.; Sen, N.; Varadi, M.; Daina, A.; Wodak, S. J.; Zoete, V.; Velankar, S.; Orengo, C. The Impact of Structural Bioinformatics Tools and Resources on SARS-CoV-2 Research and Therapeutic Strategies. Brief. Bioinform., 2021, 22 (2), 742–768. https://doi.org/10.1093/bib/bbaa362.

[15] Wise, J. Covid-19: The E484K Mutation and the Risks It Poses. BMJ, 2021, 372, n359. https://doi.org/10.1136/bmj.n359.

[16] Yurkovetskiy, L.; Wang, X.; Pascal, K. E.; Tomkins-Tinch, C.; Nyalile, T. P.; Wang, Y.; Baum, A.; Diehl, W. E.; Dauphin, A.; Carbone, C.; et al. Structural and Functional Analysis of the D614G SARS-CoV-2 Spike Protein Variant. Cell, 2020, 183 (3). https://doi.org/10.1016/j.cell.2020.09.032.

[17] Laffeber, C.; De Koning, K.; Kanaar, R.; Lebbink, J. H. Experimental Evidence for Enhanced Receptor Binding by Rapidly Spreading SARS-CoV-2 Variants. bioRxiv, 2021, 2021.02.22.432357. https://doi.org/10.1101/2021.02.22.432357.

[18] Okada, P.; Buathong, R.; Phuygun, S.; Thanadachakul, T.; Parnmen, S.; Wongboot, W.; Waicharoen, S.; Wacharapluesadee, S.; Uttayamakul, S.; Vachiraphan, A.; et al. Early Transmission Patterns of Coronavirus Disease 2019 (COVID-19) in Travellers from Wuhan to Thailand, January 2020. Eurosurveillance. European Centre for Disease Prevention and Control (ECDC) February 27, 2020. https://doi.org/10.2807/1560-7917.ES.2020.25.8.2000097.

[19] Puenpa, J.; Suwannakarn, K.; Chansaenroj, J.; Nilyanimit, P.; Yorsaeng, R.; Auphimai, C.; Kitphati, R.; Mungaomklang, A.; Kongklieng, A.; Chirathaworn, C.; et al. Molecular Epidemiology of the First Wave of Severe Acute Respiratory Syndrome Coronavirus 2 Infection in Thailand in 2020. Sci. Rep., 2020, 10 (1). https://doi.org/10.1038/s41598-020-73554-7.

[20] Buathong, R.; Chaifoo, W.; Iamsirithawon, S.; Wacharapluesadee, S.; Joyjinda, Y.; Rodpan, A.; Ampoot, W.; Putcharoen, O.; Paitoonpong, L.; Suwanpimolkul, G.; et al. Multiple Clades of SARS-CoV-2 Were Introduced to Thailand during the First Quarter of 2020. Microbiol. Immunol., 2021, 1348–0421.12883. https://doi.org/10.1111/1348-0421.12883.

[21] Joonlasak, K.; Batty, E. M.; Kochakarn, T.; Panthan, B.; Kümpornsin, K.; Jiaranai, P.; Wangwiwatsin, A.; Huang, A.; Kotanan, N.; Jaru-Ampornpan, P.; et al. Genomic Surveillance of SARS-CoV-2 in Thailand Reveals Mixed Imported Populations, a Local Lineage Expansion and a Virus with Truncated ORF7a. Virus Res., 2021, 292, 198233. https://doi.org/10.1016/j.virusres.2020.198233.

[22] Hadfield, J.; Megill, C.; Bell, S. M.; Huddleston, J.; Potter, B.; Callender, C.; Sagulenko, P.; Bedford, T.; Neher, R. A. NextStrain: Real-Time Tracking of Pathogen Evolution. Bioinformatics, 2018, 34 (23), 4121–4123. https://doi.org/10.1093/bioinformatics/bty407.

[23] Genomic epidemiology of novel coronavirus - Thailand-focused subsampling https://nextstrain.org/community/fai-k/coni/Thailand?branchLabel=aa&c=GISAID_clade&d=tree&dmax=2021-02-07&fbclid=IwAR19RF7fisEmajtTO34mlTB7pDPzpvC-adGmXyLYMkdMikA_yDTVFLvcG5A (accessed Apr 27, 2021).

[24] Kabsch, W.; Sander, C. Dictionary of Protein Secondary Structure: Pattern Recognition of Hydrogen-bonded and Geometrical Features. Biopolymers, 1983, 22 (12). https://doi.org/10.1002/bip.360221211.

[25] Rost, B.; Sander, C. Conservation and Prediction of Solvent Accesibility in Protein Families. Proteins Struct. Funct. Genet., 1994, 20 (3), 216–226.

[26] Jeffrey, G. A. An Introduction to Hydrogen Bonding; Oxford University Press, 1997.

[27] Bosshard, H. R.; Marti, D. N.; Jelesarov, I. Protein Stabilization by Salt Bridges: Concepts, Experimental Approaches and Clarification of Some Misunderstandings. J. Mol. Recognit., 2004, 17 (1), 1–16. https://doi.org/10.1002/jmr.657.

[28] Ittisoponpisan, S.; Islam, S. A.; Khanna, T.; Alhuzimi, E.; David, A.; Sternberg, M. J. E. Can Predicted Protein 3D Structures Provide Reliable Insights into Whether Missense Variants Are Disease Associated? J. Mol. Biol., 2019, 431 (11), 2197–2212. https://doi.org/10.1016/j.jmb.2019.04.009.

[29] Peter, A. S.; Roth, E.; Schulz, S. R.; Fraedrich, K.; Steinmetz, T.; Damm, D.; Hauke, M.; Richel, E.; Mueller-Schmucker, S.; Habenicht, K.; et al. A Pair of Non-Competing Neutralizing Human Monoclonal Antibodies Protecting from Disease in a SARS-CoV-2 Infection Model. bioRxiv, 2021, 2021.04.16.440101. https://doi.org/10.1101/2021.04.16.440101.

[30] Walls, A. C.; Park, Y. J.; Tortorici, M. A.; Wall, A.; McGuire, A. T.; Veesler, D. Structure, Function, and Antigenicity of the SARS-CoV-2 Spike Glycoprotein. Cell, 2020, 181 (2). https://doi.org/10.1016/j.cell.2020.02.058.

[31] Volz, E.; Hill, V.; McCrone, J. T.; Price, A.; Jorgensen, D.; O’Toole, Á.; Southgate, J.; Johnson, R.; Jackson, B.; Nascimento, F. F.; et al. Evaluating the Effects of SARS-CoV-2 Spike Mutation D614G on Transmissibility and Pathogenicity. Cell, 2021, 184 (1). https://doi.org/10.1016/j.cell.2020.11.020.

[32] Nyunt, M. H.; Soe, H. O.; Aye, K. T.; Aung, W. W.; Kyaw, Y. Y.; Kyaw, A. K.; Myat, T. W.; Latt, A. Z.; Win, M. M.; Win, A. A.; et al. Surge of Severe Acute Respiratory Syndrome Coronavirus 2 Infections Linked to Single Introduction of a Virus Strain in Myanmar, 2020. Sci. Rep., 2021, 11 (1), 1–6. https://doi.org/10.1038/s41598-021-89361-7.

[33] Mat Yassim, A. S.; Asras, M. F. F.; Gazali, A. M.; Marcial-Coba, M. S.; Zainulabid, U. A.; Ahmad, H. F. COVID-19 Outbreak in Malaysia: Decoding D614G Mutation of SARS-CoV-2 Virus Isolated from an Asymptomatic Case in Pahang. Mater. Today Proc., 2021. https://doi.org/10.1016/j.matpr.2021.02.387.

[34] Li, Q.; Wu, J.; Nie, J.; Li, X.; Huang, W.; Correspondence, Y. W.; Zhang, L.; Hao, H.; Liu, S.; Zhao, C.; et al. The Impact of Mutations in SARS-CoV-2 Spike on Viral Infectivity and Antigenicity. 2020. https://doi.org/10.1016/j.cell.2020.07.012.

[35] Shang, J.; Ye, G.; Shi, K.; Wan, Y.; Luo, C.; Aihara, H.; Geng, Q.; Auerbach, A.; Li, F. Structural Basis of Receptor Recognition by SARS-CoV-2. Nature, 2020, 581 (7807), 221–224. https://doi.org/10.1038/s41586-020-2179-y.

[36] Liu, Y.; Liu, J.; Plante, K. S.; Plante, J. A.; Xie, X.; Zhang, X.; Ku, Z.; An, Z.; Scharton, D.; Schindewolf, C.; et al. The N501Y Spike Substitution Enhances SARS-CoV-2 Transmission. bioRxiv, 2021, 2021.03.08.434499. https://doi.org/10.1101/2021.03.08.434499.

[37] Shahhosseini, N.; Babuadze, G. (Giorgi); Wong, G.; Kobinger, G. P. Mutation Signatures and In Silico Docking of Novel SARS-CoV-2 Variants of Concern. Microorganisms, 2021, 9 (5), 926. https://doi.org/10.3390/microorganisms9050926.

[38] Department of Disease Control. Thailand situation update on 19 December 2020 https://ddc.moph.go.th/viralpneumonia/eng/file/situation/situation-no349-191263.pdf (accessed Jul 16, 2021).

[39] Department of Disease Control. Thailand situation update on 24 December 2020 https://ddc.moph.go.th/viralpneumonia/eng/file/situation/situation-no350-241263.pdf (accessed Jul 16, 2021).

[40] Korber, B.; Fischer, W. M.; Gnanakaran, S.; Yoon, H.; Theiler, J.; Abfalterer, W.; Hengartner, N.; Giorgi, E. E.; Bhattacharya, T.; Foley, B.; et al. Tracking Changes in SARS-CoV-2 Spike: Evidence That D614G Increases Infectivity of the COVID-19 Virus. Cell, 2020, 182 (4). https://doi.org/10.1016/j.cell.2020.06.043.

[41] Huang, Y.; Yang, C.; Xu, X. feng; Xu, W.; Liu, S. wen. Structural and Functional Properties of SARS-CoV-2 Spike Protein: Potential Antivirus Drug Development for COVID-19. Acta Pharmacologica Sinica. Springer Nature September 1, 2020, pp 1141–1149. https://doi.org/10.1038/s41401-020-0485-4.

[42] Zhang, L.; Guo, H. Biomarkers of COVID-19 and Technologies to Combat SARS-CoV-2. Adv. Biomark. Sci. Technol., 2020, 2, 1–23. https://doi.org/10.1016/j.abst.2020.08.001.

[43] Xia, S.; Liu, M.; Wang, C.; Xu, W.; Lan, Q.; Feng, S.; Qi, F.; Bao, L.; Du, L.; Liu, S.; et al. Inhibition of SARS-CoV-2 (Previously 2019-NCoV) Infection by a Highly Potent Pan-Coronavirus Fusion Inhibitor Targeting Its Spike Protein That Harbors a High Capacity to Mediate Membrane Fusion. Cell Res., 2020, 30 (4), 343–355. https://doi.org/10.1038/s41422-020-0305-x.

[44] Sanjuán, R.; Nebot, M. R.; Chirico, N.; Mansky, L. M.; Belshaw, R. Viral Mutation Rates. J. Virol., 2010, 84 (19), 9733–9748. https://doi.org/10.1128/jvi.00694-10.

[45] David A Steinhauer, Holland, J. J. Rapid Evolution of RNA Viruses. Annu. Rev. Microbiol., 1987, 41 (1), 409–431. https://doi.org/10.1146/annurev.mi.41.100187.002205.

[46] Amraie, R.; Napoleon, M. A.; Yin, W.; Berrigan, J.; Suder, E.; Zhao, G.; Olejnik, J.; Gummuluru, S.; Muhlberger, E.; Chitalia, V.; et al. CD209L/L-SIGN and CD209/DC-SIGN Act as Receptors for SARS-CoV-2 and Are Differentially Expressed in Lung and Kidney Epithelial and Endothelial Cells. bioRxiv. bioRxiv June 23, 2020. https://doi.org/10.1101/2020.06.22.165803.

[47] Kane, L. Review 1: “KIM-1/TIM-1 Is a Receptor for SARS-CoV-2 in Lung and Kidney” License: Creative Commons Attribution 4.0 International License (CC-BY 4.0). Rapid Rev. COVID-19, 2020. https://doi.org/10.1162/2e3983f5.919b71a6.

[48] Cantuti-Castelvetri, L.; Ojha, R.; Pedro, L. D.; Djannatian, M.; Franz, J.; Kuivanen, S.; van der Meer, F.; Kallio, K.; Kaya, T.; Anastasina, M.; et al. Neuropilin-1 Facilitates SARS-CoV-2 Cell Entry and Infectivity. Science (80-.)., 2020, 370 (6518), 856–860. https://doi.org/10.1126/science.abd2985.

[49] Daly, J. L.; Simonetti, B.; Klein, K.; Chen, K. E.; Williamson, M. K.; Antón-Plágaro, C.; Shoemark, D. K.; Simón-Gracia, L.; Bauer, M.; Hollandi, R.; et al. Neuropilin-1 Is a Host Factor for SARS-CoV-2 Infection. Science (80-.)., 2020, 370 (6518), 861–865. https://doi.org/10.1126/science.abd3072.

[50] Lam, S. D.; Bordin, N.; Waman, V. P.; Scholes, H. M.; Ashford, P.; Sen, N.; van Dorp, L.; Rauer, C.; Dawson, N. L.; Pang, C. S. M.; et al. SARS-CoV-2 Spike Protein Predicted to Form Complexes with Host Receptor Protein Orthologues from a Broad Range of Mammals. Sci. Rep., 2020, 10 (1), 16471. https://doi.org/10.1038/s41598-020-71936-5.

[51] David, A.; Razali, R.; Wass, M. N.; Sternberg, M. J. E. Protein-Protein Interaction Sites Are Hot Spots for Disease-Associated Nonsynonymous SNPs. Hum. Mulat., 2012, 33 (2), 359–363. https://doi.org/10.1002/humu.21656.

[52] David, A.; Sternberg, M. J. E. The Contribution of Missense Mutations in Core and Rim Residues of Protein-Protein Interfaces to Human Disease. J. Mol. Biol., 2015, 427 (17), 2886–2898. https://doi.org/10.1016/j.jmb.2015.07.004.

[53] Jubb, H. C.; Pandurangan, A. P.; Turner, M. A.; Ochoa-Montaño, B.; Blundell, T. L.; Ascher, D. B. Mutations at Protein-Protein Interfaces: Small Changes over Big Surfaces Have Large Impacts on Human Health. Progress in Biophysics and Molecular Biology. Elsevier Ltd September 2017, pp 3–13. https://doi.org/10.1016/j.pbiomolbio.2016.10.002.

