## Supplementary Table 1 for "The SARS-CoV-2 infection in Thailand: analysis of spike variants complemented by protein structure insights"

**Supplementary Table 1: Analysis of variants found in Thailand.** The variants that were found exclusively in Thailand are indicated by *. Deleterious predictions are shown in bold.

| Variant | % | GISAID Clade ^1^ | Location  on Spike | Structural Consequences (Missense3D) | Note |
| --- | --- | --- | --- | --- | --- |
| L5F | 4.71 | GH | - | N/A (residue not contained in PDB) |  |
| S12F | 0.50 | GR | - | N/A (residue not contained in PDB) |  |
| V47A | 0.17 | GH | interface | No structural damage detected | 2 |
| T51N* | 0.17 | GH | surface | H-bond formed: Asn51 - His49 (no damage alert) | 2 |
| Q52K | 0.17 | GH | interface | No structural damage detected | 2 |
| L54F | 0.34 | G | surface | No structural damage detected | 3 |
| F55L | 0.17 | GH | buried | No structural damage detected | 2 |
| P57T* | 0.17 | GH | buried | H-bond formed: Thr57 - Gln271 (no damage alert) | 2 |
| I68R* | 0.34 | O | surface | No structural damage detected |  |
| H69Y | 0.17 | GH | surface | No structural damage detected |  |
| V70L | 0.17 | GH | - | N/A (residue not contained in PDB) | 2 |
| S71P | 0.17 | GH | - | N/A (residue not contained in PDB) | 2 |
| G75V | 0.17 | GR | - | N/A (residue not contained in PDB) |  |
| S98F | 1.51 | GR | surface | H-bond disrupted: Ser98 - Lys97 (no damage alert) |  |
| D138H | 0.34 | GH | buried | **Buried salt bridge disrupted: Asp138 - Arg21 / buried to exposed switch** |  |
| M153I | 0.50 | GH | surface | No structural damage detected |  |
| M153T | 0.34 | GR | surface | No structural damage detected |  |
| A163V | 0.17 | GH | surface | No structural damage detected |  |
| G181V | 0.17 | GH | surface | **Disallowed phi psi angles / cavity contraction / H bond formed: Ser98 - Val181** |  |
| R190S | 0.17 | GH | surface | **Cavity contraction / H-bond disrupted: Arg190 - Lys206** |  |
| R190K | 0.17 | GV | surface | **Cavity contraction / H-bond disrupted: Arg190 - Lys206** |  |
| D198G | 0.17 | GH | interface | No structural damage detected |  |
| S205T* | 0.17 | S | surface | **H-bond formed: Thr205 - Glu191 / Cavity contraction** |  |
| R214P | 0.17 | G | surface | **Disallowed phi psi angles** | 3 |
| A222V | 0.67 | GV | buried | No structural damage detected |  |
| K278T* | 0.17 | GH | surface | H-bond formed: Thr278 - Thr286 (no damage alert) |  |
| I358T | 0.17 | GH | buried | No structural damage detected |  |
| S359N | 0.17 | G | surface | H-bond formed: Asn360 - Asn359 (no damage alert) |  |
| V407A | 0.50 | S/GH/GR | surface | **6XR8 - cavity expansion**  **4M0J - no structural damage detected** | 4 |
| S459F | 20.17 | GH | surface | 6XR8 - H-bond disrupted: Arg457 - Ser459 (no damage)  4M0J - H-bond disrupted: Ser459 - Asn460 (no damage) |  |
| S477N | 1.18 | GH | surface | No structural damage detected (both 6XR8 and 4M0J) |  |
| V483F | 0.17 | GH | surface | No structural damage detected (both 6XR8 and 4M0J) |  |
| E484D | 0.17 | GR | surface | No structural damage detected (both 6XR8 and 4M0J) |  |
| N501T | 1.51 | GR | interface (ACE) | 6XR8 and 4M0J - H-bond disrupted: Asn501- Gly496 and Asn501 - Tyr505, H-bond formed: Thr501- Gln498 (no damage alert)  4M0J only - H-bond formed: Thr501 - Gly502 (no damage alert) |  |
| K537R | 0.17 | GH | surface | H-bond formed: Arg537 - Asn536 (no damage alert) |  |
| D614G | 48.74 | G/GH/GR/ GRY/GV | interface | **H-bond disrupted: Asp614 (chain A) - Lys835 (chain B) and Asp614 (chain A) - Lys854 (chain B) / buried charge replaced** |  |
| Q675H | 0.17 | GR | surface | No structural damage detected |  |
| A688P | 0.17 | GH | - | N/A (residue not contained in PDB) | 2 |
| V785L | 0.50 | GV | buried | No structural damage detected |  |
| T791I | 0.17 | GR | surface | **Cavity contraction / H-bond disrupted: Thr791 - Pro792** |  |
| A829T | 39.16 | S | surface | H-bond formed: Thr829 - Asp830 (no damage alert) |  |
| G832C* | 0.34 | G | interface | No structural damage detected |  |
| F855L | 0.17 | GR | interface | No structural damage detected |  |
| D936Y | 0.50 | GH | surface | No structural damage detected |  |
| S939F | 2.02 | GH | surface | No structural damage detected |  |
| D1118Y | 0.34 | GH | surface | **Cavity contraction** |  |
| S1147L | 0.17 | GH | surface | H-bond disrupted: Ser1147 - Pro1143 (no damage alert) |  |
| P1162L | 0.17 | S | surface | No structural damage detected |  |
| V1176F | 0.17 | GR | - | N/A (residue not contained in PDB) |  |
| E1202Q | 0.34 | GV/GH | - | N/A (residue not contained in PDB) |  |
| G1219C | 0.50 | L | - | N/A (residue not contained in PDB) |  |
| D1260Y | 0.17 | GR | - | N/A (residue not contained in PDB) | 5 |
| Note: 1) The column shows on the GISAID clade of strains in which the majority of variants can be found. Some variants are not directly mappable onto the phylogenetic tree due to 2) variant from GISAID ID: 708814, 3) incomplete genome information, 4) one record from GISAID ID 708814, one record had incomplete genome, one record had incomplete collection date, and 5) incomplete date of collection. | | | | | |
