## Supplementary File 1-4 for "The SARS-CoV-2 infection in Thailand: analysis of spike variants complemented by protein structure insights"

**Supplementary File 1 – analysis of surface-interface residues on PDB coordinate 6XR8 and 6M0J**

(Supplementary File 1 - Surface-interface.xlsx)

<https://drive.google.com/file/d/1-eHYWmcAQ5yo2S2JhhPhjYnIEvGE8PVn/view?usp=sharing>

**Supplementary File 2 – variant distribution**

(Supplementary File 2 - Variant distribution.xlsx)

<https://drive.google.com/file/d/1-fJgPKKHZiGm8Y1-n9FKy7cnn31qbRx2/view?usp=sharing>

**Supplementary File 3 - distribution of variants on the spike protein complex**

(Supplementary File 3 - Variant distribution - All.mp4)

<https://drive.google.com/file/d/12p1ZV_poR8okwOtPDJZ0U46y5Zv2XgWe/view?usp=sharing>

**Supplementary File 4 - distribution of Thai variants on the spike protein complex**

(Supplementary File 4 - Variant distribution - Thai.mp4)

<https://drive.google.com/file/d/12oVYg6ZBQ-mUi_nvWbvP55RL6ffLOrbS/view?usp=sharing>
