## Supplementary Figure 1 for "The SARS-CoV-2 infection in Thailand: analysis of spike variants complemented by protein structure insights"

**Appendix: Supplementary Materials**

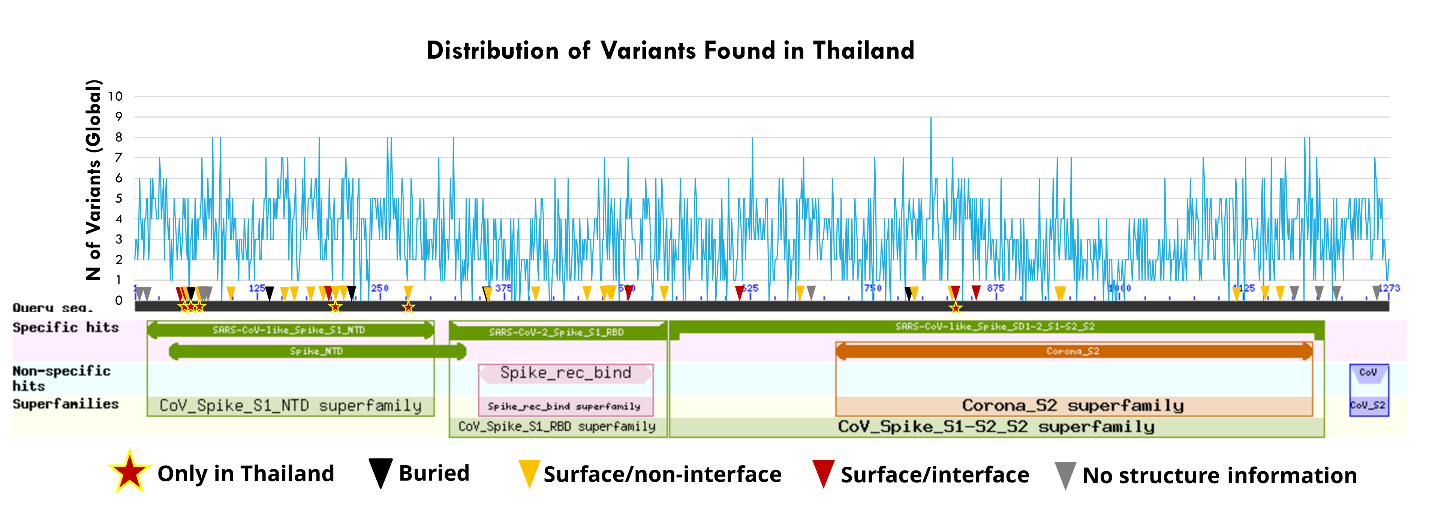
 **Supplementary Figure 1: Distribution of variants on the spike protein.** Residues where Thai variants were found are indicated by arrowheads (black: buried, yellow: surface/non-interface, red: surface/interface, grey: no structure information). Variants specific to Thailand are indicated by ★. Colour boxes depict spike conserved domains.
